# Post-transcriptional Regulation Coordinating Transcription and Translation During Circadian Oscillation and Stress Recovery in Plants

**DOI:** 10.1101/2025.09.04.674151

**Authors:** Pengfei Xu, You Wu, Qihui Wan, Xiang Yu

## Abstract

Circadian rhythms orchestrate gene expression to align plant growth and development with daily environmental cycles. However, the post-transcriptional mechanisms that coordinate transcriptional and translational rhythmicity remain incompletely understood. To address this, we analyzed time-series transcriptome and translatome profiles in Arabidopsis seedlings, identifying 5,185 genes with rhythmicity at both levels. These genes were classified into four distinct groups based on phase and amplitude differences between transcription and translation. Circadian mRNAs with high oscillation amplitudes tended to undergo co-translational RNA decay (CTRD), whereas intronless genes displayed the lowest amplitudes, likely due to their mRNA instability and short half-lives. While CTRD and NAD⁺ capping modulate amplitude differences, intronless and circadian translational efficiency (TE) influence both phase and amplitude variations. Additionally, CTRD, NAD^+^ capping and circadian TE facilitate fast recovery of heat-induced genes to normal hemostasias. Collectively, our findings demonstrate that these post-transcriptional regulation shapes both synchronized and decoupled transcription and translation during plants response to diel and environmental dynamics.

## INTRODUCTION

Circadian rhythms orchestrate the fine-tuning of physiological and molecular events, including metabolism, growth, behavior, and evolutionary adaptation to daily and seasonal environmental changes (*1*). Circadian gene oscillations are driven either by an internal clock or by external cues, which include various light signals and temperature fluctuations (*2*). Transcription has long been regarded as the main contributor to mRNA abundance. The morning-phased transcription factors CCA1 and LHY inhibit the expression of afternoon-phased *PRR1*, *PRR3*, *PRR5*, *PRR7*, and *PRR9*, as well as evening-phased *ELF3*, *ELF4*, and *LUX*. Conversely, the PRRs suppress the transcription of *CCA1* and *LHY*, forming a regulatory feedback loop (*3*). The ELF3-ELF4-LUX evening complex negatively regulates the transcription of PRR7 and PRR9 (*4*). *TOC1* is sequentially expressed throughout the daytime and represses the transcription of *CCA1* and *LHY* (*5*). Thousands of genes have been found to exhibit circadian oscillations over a 24-hour period (*6*). Those genes involved in plant development, metabolic and signaling pathways, response to abiotic and biotic stress (*2, 7, 8*). For instance, *RVEs* genes have been found to function as core circadian clock activators, modulating the regulation of ELF3-ELF4-LUX evening complex, *TOC1* and *PRR5* expression (*9*). When Arabidopsis is subjected to heat stress at 37°C, the circadian clock proteins RVE4 and RVE8 have been reported to initiate heat shock (HS)-inducible gene expression independently of heat shock factors (HSFs). These proteins partially regulate plant thermotolerance by modulating the expression of downstream transcription factors *ERF53* and *ERF54*, specifically around noon. This finding indicates that the circadian clock can control the magnitude of transcript abundance changes depending on the time of day (*10*). Moreover, numerous key circadian genes have been demonstrated to regulate flowering time in plants, a process crucial for their adaptation to varying latitudinal environments (*11–13*).

Several studies have shown that the circadian oscillators also undergo epigenetic modifications and post-transcriptional regulation. DNA methylation, and alternative splicing of pre-mRNA affect the expression of circadian clock genes. Epigenetic modifications of *CCA1* and *LHY*, along with their reciprocal regulators *TOC1* and *GI*, mediate changes in the expression of downstream genes and pathways (*14*). Analysis of alternative splicing patterns in Arabidopsis cultured under constant light and temperature conditions uncovered hundreds of mRNAs regulated by the circadian clock, displaying circadian rhythms in their alternative splicing (*15*). Notably, several studies on diurnal gene expression have revealed discrepancies between mRNA and protein abundance, highlighting the impact of post-transcriptional regulation, including translational control. *PRR9*, *PRR7*, *PRR5*, and *TOC1* are expressed sequentially throughout the daytime, with the peak protein levels lagging behind peak transcript levels by 4-8 h (*16*). A study integrating transcriptome and translatome analyses revealed that 36% of mRNAs exhibited rhythmicity exclusively at the translational level under circadian conditions (*17*). Another study revealed that translational regulation mediated by upstream open reading frames (uORFs) in the 5′UTR contributes to the diel oscillation of mORF translation (*18*). These studies indicate that translational regulation plays a substantial role in shaping circadian expression patterns.

Accumulating evidence suggests that post-transcriptional regulation plays significant roles in gene oscillation (*19–22*). Many genes that alter clock phase when mutated affect both RNA splicing and RNA decay. Mutation of *LSM4* and *LSM5*, which are components of a core spliceosome, causes a long-period phenotype (*23*). Co-translational RNA decay (CTRD), a process in which are turned over by 5’-3’ exonuclease while still associated with translating ribosome, was first uncovered in yeast (*24*), and has subsequently been widely observed in plants (*25, 26*). Diurnal mRNAs with large amplitudes tend to be CTRD in tomato, suggesting that CTRD contributes to the rapid turnover of diurnal mRNAs (*27*). However, the function of RNA stability in regulating the amplitude changes remains largely unknown.

When external signals overload the ambient limit in plants, the circadian oscillators help organisms to anticipates the stress period and prepares the plant to fight against it (*2*). It remains largely unknown how post-transcriptional regulation assists plants in rapidly adapting to transient extreme stress events during the day, and how it facilitates recovery to normal physiological states after stress. Previous studies have shown that plants can generally recover to their initial state within hours after transient stress exposure; however, the role of post-transcriptional regulation in this recovery process remains largely unclear.

Through time-course transcriptome and translatome profiling, we investigated how post-transcriptional regulations contribute to coordinating circadian oscillations of transcriptional and translation. Our study revealed that the features of post-transcriptional regulations, CTRD, intronless, NAD⁺capping and circadian translational efficiency (TE), influence the phase and amplitude differences between transcription and translation. Notably, the impact of CTRD on large-amplitude oscillations is conserved across plants, whereas intronless genes predominantly contribute to low-amplitude oscillations of diel-regulated genes. Additionally, CTRD, NAD⁺ capping, and circadian TE contributed to the fast recovery of both transcription and translation after heat stress. Together, our findings demonstrate that post-transcriptional regulation shapes both synchronized and asynchronous transcription and translation in plants, providing new insights into the regulation of circadian gene oscillations and resilience to environmental stress.

## RESULTS

### Overview of the synchronous and asynchronous rhythmicity of circadian transcription and translation in plants

To investigate the coordination of circadian rhythmicity of genes at transcriptional and translational level, we reanalyzed the time-series transcriptome and translatome from wild-type Arabidopsis (Col-0) seedlings using the MetaCycle2 R package (*28*). The seedings were collected every 3 hours over a full 24-hour time course (following 10 days of entrainment and then 2 days in free-running conditions. ZT48: beginning of the light period; ZT72: end of the night) (*17*) (Figure 1A). In total, we identified 6,084 and 9,724 genes exhibiting significant circadian oscillations at the transcriptome and translatome levels, respectively (p < 0.01) (Table S1). A greater number of genes displaying oscillation at translational level, reflecting the heterogeneity between transcriptional and translational regulation. Among them, the 5,185 genes displaying circadian oscillations at both levels (Figure 1B). These genes were enriched in circadian processes, responses to stress stimulus, cellular processes, and photosynthesis (Figure S1).

**Figure 1.**
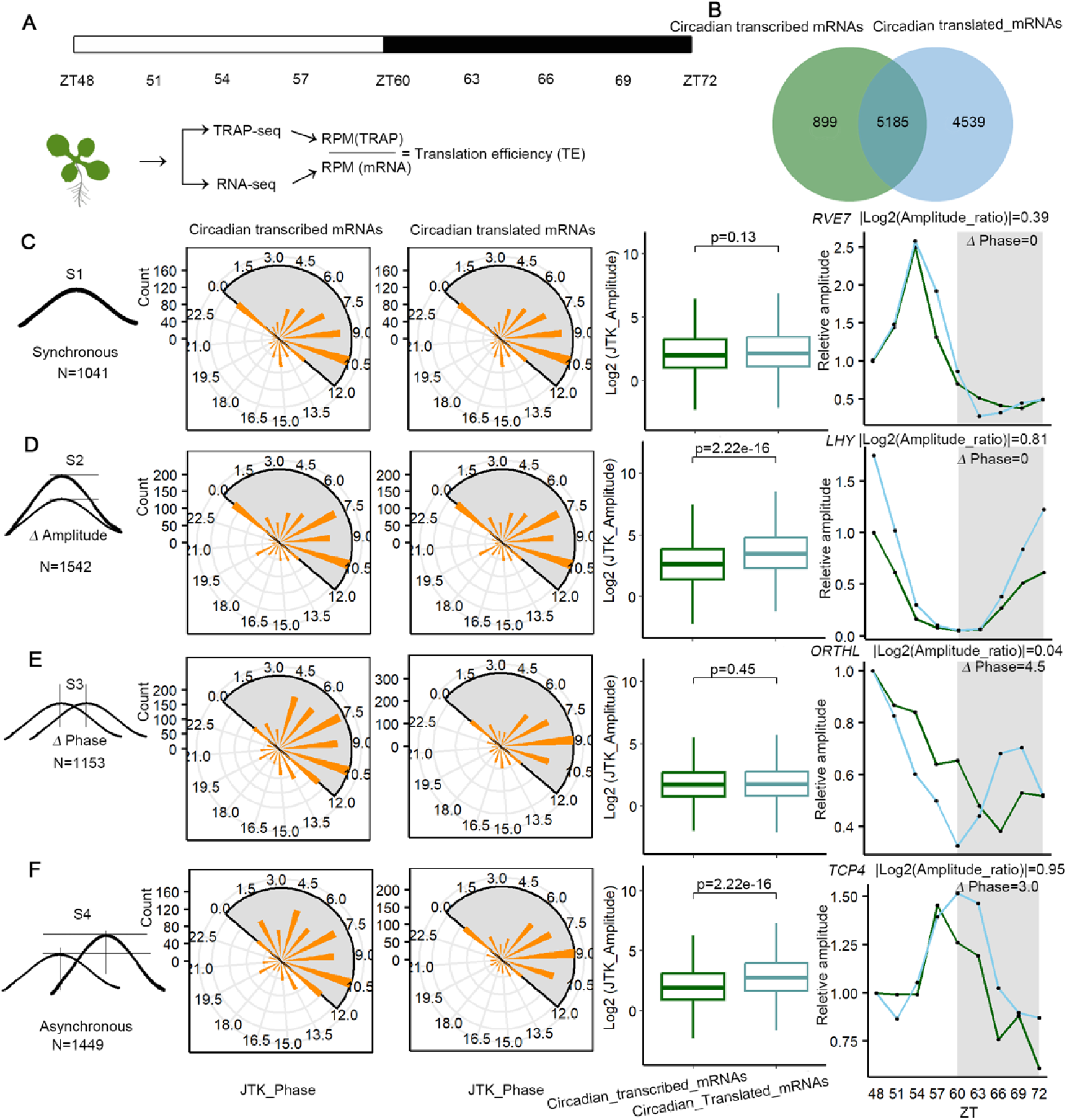
Analysis of the synchronous and asynchronous rhythmicity of genes with circadian transcription and translation in plants. (A) Schematic overview of the experimental design. After 10 days of entrainment under light/dark cycles, seedlings were transferred to constant light and temperature conditions (22 °C) for 2 days (days 11–12). Samples were collected for mRNA-Seq and TRAP-Seq analyses (*17*). (B) Venn diagram showing the overlap of circadian transcribed mRNAs and circadian translated mRNAs. (C-F) Classification of four groups based on differences in their phase and amplitude patterns. From left to right: panel 1: Diagram depicting the relative phase positions and amplitude differences between rhythmically circadian mRNAs and circadian translated mRNAs; panel 2-3: The circle plot Circular phase distribution plots of circadian transcribed mRNAs (panel 2) and circadian translated mRNAs (panel 3). The Value indicates the gene number; panel 4: Boxplot comparison of oscillation amplitudes between circadian transcribed mRNAs and circadian translated mRNAs; panel 5: Representative line plots showing relative amplitude patterns of selected genes at both transcriptional (green lines) and translational (blue lines) levels.

Next, we performed an intersection analysis of the 5,185 circadian mRNAs to further uncover the heterogeneity between genes exhibiting rhythmic transcription and translation, which allowed us to classify the genes into four groups based on differences in their phase and amplitude patterns. The classification criteria for S1-S4 groups were defined as follows: S1 group: |Δphase| < 1.5 h and |Log_2_(amplitude fold change) | < 0.58; S2 group: |Δphase| < 1.5 h and |Log_2_(amplitude fold change) | ≥ 0.58; S3 group: |Δphase| ≥ 1.5 h and |Log_2_(amplitude fold change) | < 0.58; S4 group: |Δphase| ≥ 1.5 h and |Log_2_(amplitude fold change) | ≥ 0.58 (Figure 1C-F). This analysis revealed that S1 group consisted of 1,041 genes with synchronous transcriptional and translational rhythms, suggesting that these genes were primarily regulated at transcription level. The S2 group included 1,542 genes that displayed similar phases but differed in amplitude (Figure 1D). Analysis of circadian phases revealed that total mRNA abundance peaked primarily at ZT7.5 and ZT10.5, whereas translated mRNAs exhibited a similar phase distribution. As expectation, there was a significant difference in amplitude was observed between transcribed and translated mRNAs (Figure 1D). S3 group exhibited similar amplitudes but showed substantial differences in phase (Figure 1E). Analysis of the circadian phases of these genes showed that transcribed mRNAs abundance tended to peak at ZT7.5 and ZT10.5, whereas translated mRNAs abundance peaked around ZT9.5. Boxplot analysis of amplitudes indicated no significant differences between transcribed mRNAs and translated mRNAs abundance. The S4 group contained 1,449 genes that differed in both phase and amplitude indicating that these genes are subject to distinct rhythmic regulatory mechanisms at the transcriptional and translational levels (Figure 1F). Analysis of the circadian phases and amplitudes of these genes revealed distinct differences in both phase distribution and oscillation amplitude.

For instance, the core circadian gene *LHY* in S2 showed a higher amplitude at the translational level. *ORTHL* in S3 had a |log_2_(amplitude fold change)| of 0.04 and a phase shift of 4.5 h, with translated mRNAs peaking earlier than transcribed mRNAs (Figure 1E). *TCP4* in group S4 showed a 3 h phase advance in transcribed mRNAs compared to translated mRNAs, while translated mRNAs amplitude was higher than that of transcribed mRNAs (Figure 1F). Taken together, the genes in group S2, S3 and S4 exhibited asynchronous phase and/or amplitude between transcription and translation.

### The impact of co-translational RNA decay on asynchronous circadian rhythms of transcription and translation

To explore the potential mechanism of asynchronous rhythm between transcription and translation, we analyzed the effect of post-transcriptional regulation, including the features of intronless without RNA splicing, miRNA-targeted degradation, CTRD, and NAD^+^ capping (a non-canonical 5′ cap structure, the NAD⁺ cap). Genes with these features tend to be less stable than other mRNAs in plants (*19, 32–35*). Firstly, we investigated the effect of CTRD, a process in which 5’-3’ exonuclease while still associated with translating ribosomes, is a widespread mechanism in plants (*26, 29*). CTRD plays a key role in the rapid turnover of diurnally regulated mRNAs (*27*). Based on this observation, we investigated whether CTRD contributes to the phase and amplitude patterns observed in circadian transcription and translation. We compiled a set of 3,635 CTRD genes (Table S2) from three previously published studies, representing a conserved group of CTRD genes in Arabidopsis (*26, 30, 31*). An intersection analysis between the CTRD genes and the 5,185 common circadian oscillating genes identified 1,262 CTRD genes exhibiting rhythmic patterns at both transcriptional and translational levels (Figure 2A). These findings suggested that co-translational RNA decay partially contributes to the regulation of circadian rhythms at both transcriptional and translational layers. We next investigated the association between CTRD and the four groups with distinct phase/amplitude expression patterns. This analysis revealed that CTRD genes were significantly overrepresented in the S2 groups, but not in the S1, S3 and S4 groups (Figure 2B), suggesting that CTRD appeared to play a predominant role in modulating the amplitude difference between rhythmic transcription and translation.

**Figure 2.**
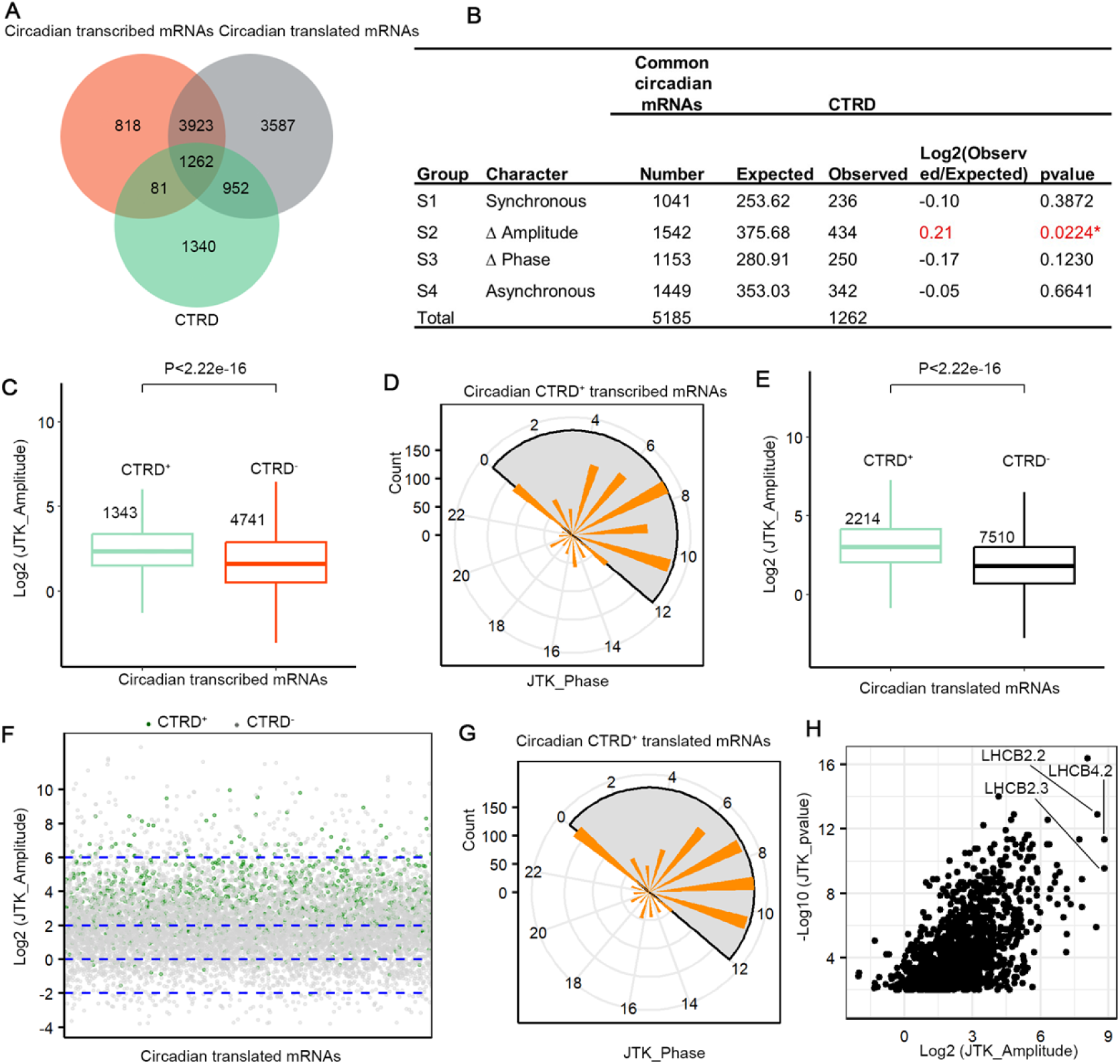
Co-translational RNA decay contributes to the high amplitude of circadian oscillations in Arabidopsis. (A) Venn diagram shows the overlap among circadian transcribed mRNAs, circadian translated mRNAs, and mRNAs undergoing CTRD. (B) genes with CTRD significantly enriched in S2 group. The ratio of expected numbers and observed numbers of genes with CTRD were calculated. Significantly over-or underrepresented groups are highlighted in red (Chi-squared test, * p<0.05, ** p<0.01). (C, E) Boxplots displaying significantly higher amplitudes of circadian transcribed mRNAs (C) and translated mRNAs (E) for genes with CTRD compared to those without CTRD. Statistical significance was assessed using the Wilcoxon test. (D, G) Phase distribution plot of circadian transcribed mRNAs (C) and translated mRNAs (G) that undergo CTRD. The y-axis indicates the number of genes. (F) Distribution of circadian translated mRNAs with (green) and without CTRD (gray) across different amplitude levels. Circadian translated mRNAs with higher amplitudes are more likely to undergo CTRD. (H) circadian translated mRNAs are enriched in photosynthesis pathways and exhibit the highest amplitude oscillations.

### Co-translational RNA decay contributed to circadian oscillations with large amplitudes in plants

To further explore the role of CTRD in circadian transcriptional and translational oscillations, we analyzed a set of 3,635 CTRD genes in Arabidopsis by intersecting them with circadian transcribed and translated mRNAs. We identified 1,343 CTRD genes overlapping with circadian transcribed mRNAs and 2,214 overlapping with circadian translated mRNAs (Figure 2A). Circadian transcribed mRNAs associated with CTRD (CTRD^+^) exhibited significantly larger oscillation amplitudes than those without CTRD (CTRD^-^) (Figure 2C). In our previous study, we found that CTRD contributed to the turnover of mRNAs with larger oscillation amplitudes in tomato (*27*). This finding suggests that CTRD-mediated regulation of circadian genes with high-amplitude is a conserved mechanism in plants. These CTRD-associated circadian mRNAs peaked predominantly at ZT7.5 and ZT10.5 (Figure 2D). We further examined the amplitude of translational profiles and found that translated mRNAs with CTRD also displayed significantly higher amplitudes than those without CTRD (Figures 2E and F). These translated genes undergoing CTRD were most highly enriched at ZT7.5, ZT9, and ZT10.5 (Figure 2G). Among the top circadian translated mRNAs with the highest amplitudes and associated with CTRD, many were involved in photosynthesis (Figure 2H). These genes exhibited rhythmic oscillations at both the transcribed mRNA and translated mRNAs. For instance, *LHCB4.2*, *LHCB3.2*, *LHCB2.2*, and *CCH1* exhibited high mRNA abundance at both transcriptional and translational levels during the light period, followed by rapid downregulation at night. This resulted in exceptionally large amplitude changes (Figure S2). Together, these findings support the role of CTRD in regulating high amplitude rhythmic oscillations not only at total mRNAs but also at actively translated mRNAs.

### Intronless transcripts exhibit low-amplitude circadian expression patterns

We found that intronless genes are the most abundant in the genomes of Arabidopsis, tomato, and rice (Figure S3A). Recently, a survey of RNA half-lives was conducted in wild-type Col-0. Based on this dataset, we firstly compared the RNA half-lives of intronless genes with those of genes containing introns in shoot and root tissues (*31*). The results showed that intronless genes have the shortest RNA half-lives compared to genes containing introns (Figure 3A and Figure S3B). We then assessed the impact of intron number on circadian gene oscillations, and found that intronless genes exhibited the lower oscillation amplitudes compared to circadian genes containing introns (Figure 3B). The short half-life of intronless genes may prevent sustained accumulation of highly abundant mRNAs, thereby contributing to their low oscillation amplitudes. To test whether the association between intronless genes and lower diurnal amplitude is conserved in plants, we analyzed two additional datasets from tomato and rice (*27, 36*). Interestingly, our analysis showed that intronless genes exhibited the lowest oscillation amplitudes in both tomato and rice datasets (Figure 3C, D and Figure S3C). Correspondingly, intronless genes in tomato displayed highest value of proportion uncapped (the ratio of normalized degraded RNA reads count compared to normalized RNA-seq reads count) representing mRNA instability (Figure 3C). Additionally, intronless genes exhibited the lower oscillation amplitudes in translated mRNAs compared to circadian genes containing introns (Figure S3D). Taken together, intronless genes exhibit short half-lives and low mRNA stability, resulting in the lowest transcriptional and translational oscillation amplitudes observed in plants. This feature may allow plants to fine-tune the expression of specific genes with minimal energetic cost, particularly in pathways that require rapid but low amplitude responses to daily environmental cycles.

**Figure 3.**
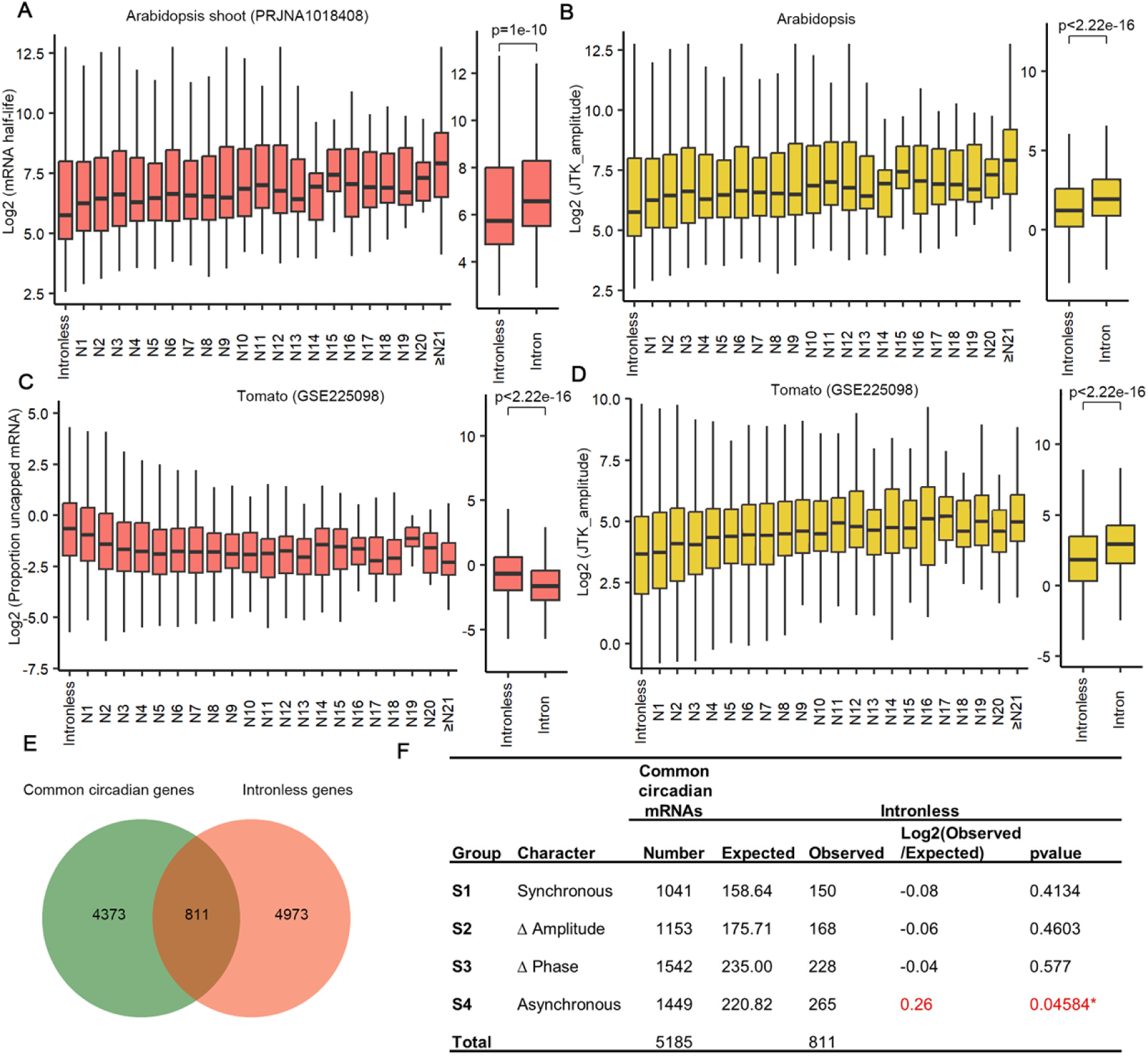
Intronless genes exhibit low circadian amplitude in plants. (A) Coding genes without introns (intronless genes) display significantly shorter half-life compared to genes with introns in Arabidopsis. (B) Intronless genes display significantly lower circadian amplitudes compared to genes with introns in Arabidopsis. (C) Coding genes without introns (intronless genes) display significantly higher proportion uncapped, indicating less stable, compared to genes with introns in Tomato. (D) Intronless genes display significantly lower circadian amplitudes compared to genes with introns tomato. (E) Venn diagram showing the overlap among common circadian genes and intronless genes. (F) Intronless genes are significantly enriched in S4 group. Expected and observed numbers of intronless genes in each group were calculated. Significantly over- or underrepresented groups are shown (Chi-squared test, * p<0.05, ** p<0.01).

Furthermore, the intersection analysis between the intronless genes and the 5,185 common circadian genes identified 811 genes exhibiting rhythmic patterns at both transcriptional and translational levels (Figure 3E). This finding suggested that intronless feature contributes to the regulation of circadian rhythms at both transcriptional and translational processes. We next investigated the association between intronless and the four groups with distinct phase/amplitude expression patterns. This analysis revealed that intronless genes were significantly overrepresented in the S4 group (Figure 3F), suggesting that the absence of introns contributes to the heterogeneity between transcribed and translated mRNAs in both amplitude and phase.

### NAD^+^-capped transcripts exhibit intermediate amplitude circadian rhythms

NAD^+^-capped transcripts have been widely identified in plant RNAs and is known to promote transcripts degradation (*21, 22*). NAD^+^-capped transcripts could also be targeted by CTRD (*35*). To examine whether NAD⁺-capped mRNAs contribute to changes in oscillation amplitude, we analyzed 5,642 NAD⁺-capped RNAs obtained from published data by comparing them with common circadian mRNAs (*22*). Among the 5,185 circadian genes, 1,707 were identified as NAD⁺-capped mRNAs (NAD^+^), while the remaining 3,478 were not (NAD^-^) (Figure 4A). Among the 1,707 genes, 573 were significantly overrepresented in S2, while 295 were underrepresented in S1. These results suggest that NAD^⁺^ capping tends to play a predominant role in modulating amplitude rather than phase. We then analyzed the amplitude differences between NAD⁺ and NAD^-^ groups and found that NAD⁺-capped circadian mRNAs exhibited significantly larger oscillation amplitudes than NAD^-^-capped mRNAs (Figure 4C). A similar trend was observed at the translational level, where NAD⁺-capped translated mRNAs also showed significantly higher amplitudes than those NAD^-^-capped mRNAs (Figure 4D). This result suggested that NAD^+^ capping also contributed to turnover of mRNAs with large oscillation amplitudes in Arabidopsis.

**Figure 4.**
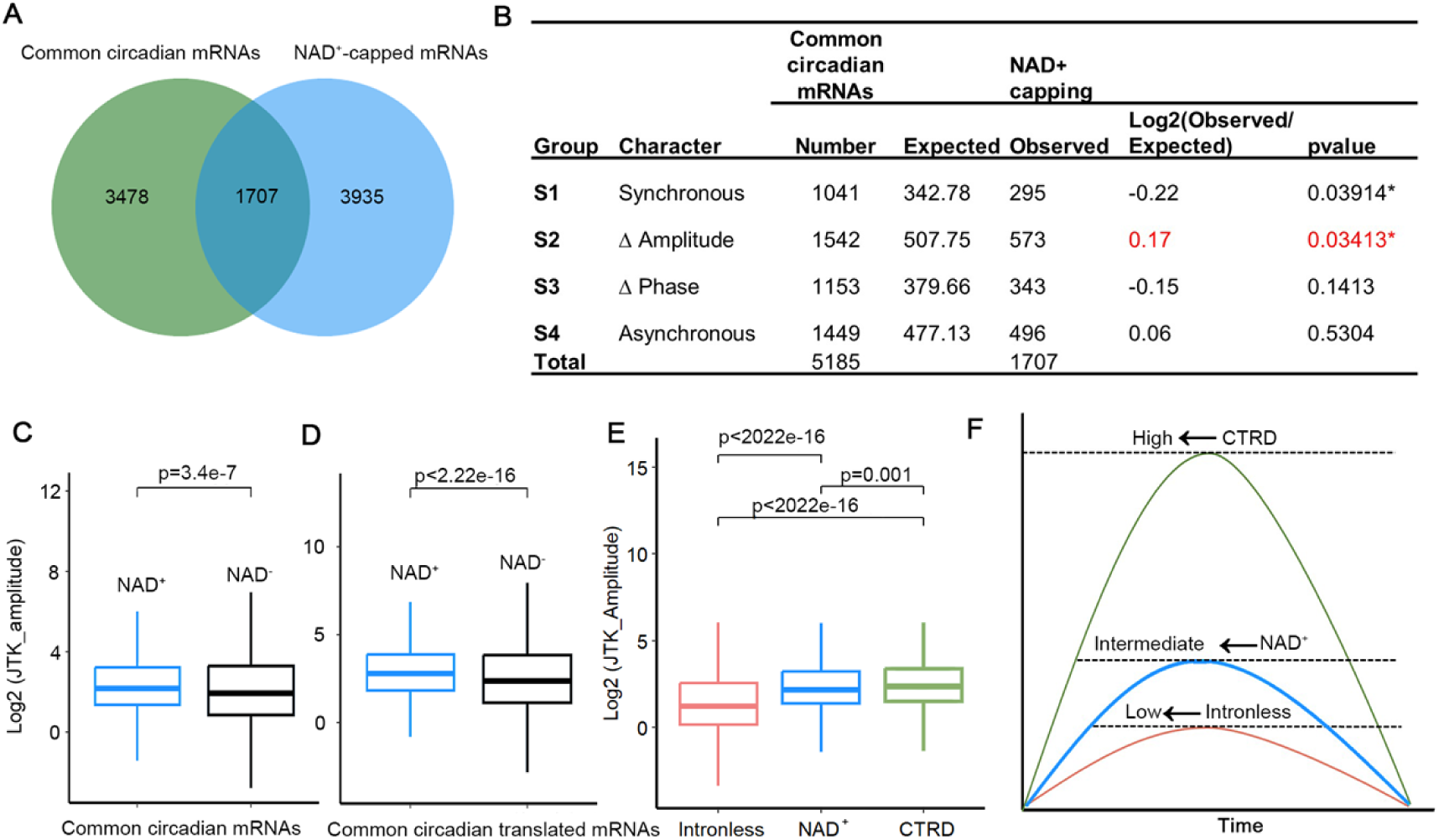
NAD+ capping contributes to the intermediate amplitude of circadian oscillations in Arabidopsis. (A) Venn diagram shows the overlap among common circadian mRNAs and NAD+-capped mRNAs. (B) NAD+ capping significantly enriched in S2 group (Chi-squared test, * p<0.05, ** p<0.01). (C, D) NAD⁺ capping contributes to the amplitude modulation of circadian transcribed mRNAs (C) and translated mRNAs (D). (E-F) Boxplot analysis (E) and schematic model (F) shows significant amplitude differences among genes with different features: CTRD-associated genes show the strongest oscillations (highest amplitudes), followed by NAD⁺-capped RNAs (intermediate), while intronless genes exhibit the weakest oscillations (lowest amplitudes).

In our previous work, we showed that miRNA cleavage efficiency oscillates in a diurnal manner in tomato (*27*). However, it remained unclear whether miRNA-mediated mRNA decay contributes to changes in mRNAs oscillation. To address this, we analyzed a dataset containing 145 experimentally verified miRNA target genes in Arabidopsis (*37*). Among these, only 34 genes exhibited circadian oscillations (Figure S4A). When comparing the amplitudes of these 34 miRNA-targeted circadian genes with other circadian genes, we found no significant difference (Figure S4B).

We next performed a comparative analysis to evaluate the effects of CTRD, intronless, and NAD⁺ capping on oscillation amplitude. The results revealed that CTRD-associated genes exhibited the highest amplitudes, consistent with their rapid transcript turnover enhancing rhythmic fluctuations. In contrast, intronless genes displayed the lowest amplitudes, likely due to their inherently reduced mRNA stability. Notably, NAD⁺-capped transcripts showed intermediate amplitude levels, suggesting a moderate contribution of NAD⁺-mediated decay to oscillation dynamics (Figure 4E and F). These findings revealed that CTRD, NAD^+^ capping and intronless status synergistically regulate the amplitude of circadian mRNA oscillations.

### The impact of translation efficiency on circadian translational rhythm

To investigate how translation efficiency (TE) affects circadian translational rhythm, we firstly performed a transcriptome-wide analysis using TRAP-seq and RNA-seq datasets. For each gene, the expression level was normalized to reads per million (RPM), and TE was calculated as the ratio of TRAP-seq RPM to RNA-seq RPM (Figure 1A). Time-series TE data were analyzed using the MetaCycle R package, identified 379 genes with significant circadian oscillations in TE (circadian TE) (p < 0.01), suggesting that the TE of these genes was regulated by circadian rhythm (Figure 5A). The circadian TE showed predominant peaks at ZT 0 and ZT 1.5 (Figure 5B). To further explore global expression patterns, we used the Mfuzz R package to perform unsupervised clustering of TE profiles, grouping them into four distinct clusters (Figure 5C). Cluster 1 included 89 genes with highest TE at ZT 6 or ZT 9, Cluster 2 had 110 genes with lowest TE at ZT 12, Cluster 3 comprised 73 genes with highest TE at ZT 15, and Cluster 4 contained 107 genes with lowest TE at ZT 6, ZT 9 or ZT 12 (Figure 5C). This unsupervised clustering provided a comprehensive overview of the oscillatory dynamics of TE in Arabidopsis.

**Figure 5.**
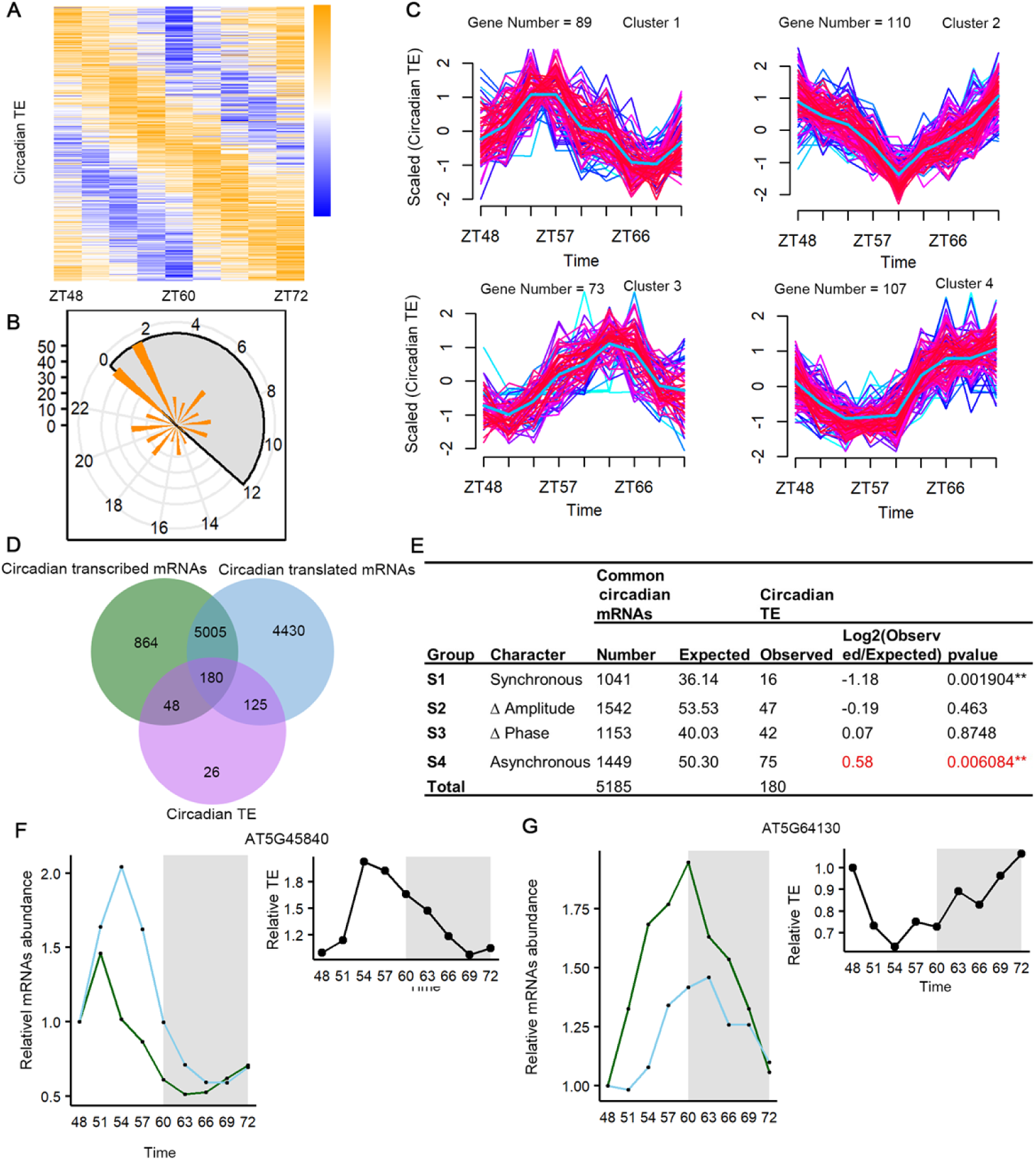
The impact of circadian translation efficiency on asynchronous transcriptional and translational rhythmicity. (A) The heatmap shows genes with relative circadian TE across the 9 time points identified by R Metacycle package. (B) The distribution of genes with circadian TE in different phases. (C) Distribution of genes with scaled circadian TE pattern in 4 clustered groups. (D) Venn diagram shows the overlap among circadian transcribed mRNAs, circadian translated mRNAs and genes with circadian TE. (E) genes Circadian TE are significantly enriched in S4 groups (Chi-squared test, * p<0.05, ** p<0.01). (F, G) Two representative genes exhibiting relative transcribed mRNA abundance (green line, left), translated mRNA abundance (blue lines, left), and relative TE (black line, right) of *AT5G45840* (F) and *AT5G64130* (G).

Intersection analysis of circadian transcribed mRNAs, circadian translated mRNAs, and rhythmic TE identified 180 genes exhibiting rhythmic patterns at all three levels (Figure 5D). We next examined the association between circadian TE and the four groups with distinct phase/amplitude expression patterns. This analysis revealed that circadian TE was significantly overrepresented in the S4 group (Figure 5E), suggesting that TE rhythmicity contributes to phase and amplitude differences between transcribed and translated mRNAs. For instance, *AT5G45840* in S4 group, a leucine-rich-repeat receptor-like kinase, showed a 3 h phase advance in transcribed mRNAs compared to translated mRNAs, while translated mRNAs amplitude was significantly higher than that of transcribed mRNAs. Moreover, its TE was also exhibited circadian oscillation (Figure 5F). Similarly, *AT5G64130* with a circadian oscillation of TE, encoding a cAMP-regulated phosphoprotein 19-related protein, displayed earlier peaks in transcribed mRNAs compared to translated mRNAs, along with higher translated mRNA amplitude and circadian TE oscillation (Figure 5G).

### Plants utilize multiple post-transcriptional mechanisms to recover transcription and translation homeostasis after transient heat stress

Although post-transcriptional regulation plays an important role in maintaining intrinsic rhythmic oscillations, it remains unclear how it rapidly assists plants in returning to normal states after experiencing transient heat stress. In this study, we reanalyzed the datasets sequenced from 4 time points. After being 1 hour 37°C heat stress, the samples were collected at ZT54 (0 h of recovery) and at 1, 3, and 6 h of recovery (ZT55, ZT57 and ZT60), respectively (Figure 6A) (*17*). In parallel, a set of samples grown under normal conditions at 22 °C was used as the control group. To assess the transcriptome-wide impact of transient heat stress, we first performed principal component analysis (PCA). After 1 hour of heat stress at 37 °C (Recovery 0h), both transcribed mRNAs and translated mRNAs profiles showed clear structural separation from the control group, indicating a strong global response (Figure S5A and B). The heat stress had a pronounced effect on gene expression, resulting in a significant outlier effect across the transcriptome and translatome. After 1 hour of recovery at 22 °C, the outlier effect began to diminish (Figure S5A and B). By 3 hours of recovery, PCA revealed that the treatment and control groups had started to converge, making their separation less distinct (Figure S5A and 5B).

**Figure 6.**
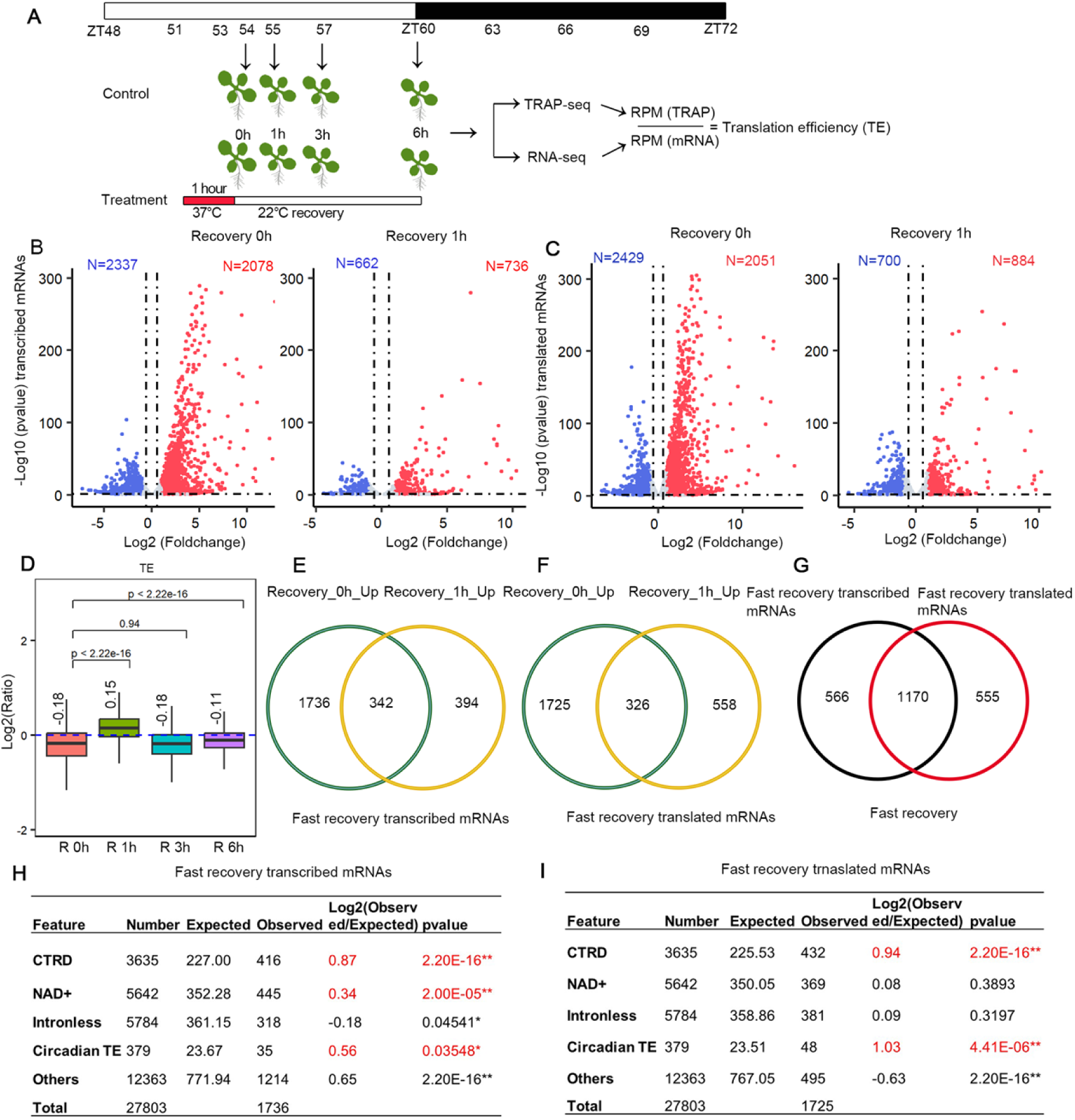
Plants coordinate multiple post-transcriptional mechanisms to restore highly accumulated transcribed and translated mRNAs induced by heat stress to normal hemostasias. (A) Schematic overview of the heat stress experiment. Samples were collected at ZT54 (immediately after heat stress, defined as 0 h of recovery), and after 1, 3, and 6 hours of recovery, corresponding to ZT55, ZT57, and ZT60, respectively. (B) Volcano plot shows differentially expressed genes (DEGs) after 1 hour of heat stress at 37 °C (left), and DEGs followed by a subsequent 1h recovery period. (C) Volcano plot shows differentially translated genes (DTGs) after 1 hour of heat stress at 37 °C (left), and DTGs followed by a subsequent 1h recovery period. (D) The foldchange of TE after 1 hour of heat stress and different recovery periods (1 h, 3 h and 6 h) compared to the control samples without heat stress. (E-F) Venn diagram analysis identifies 1736 heat-induced genes as fast-recovery transcribed mRNAs that rapidly restore to normal mRNA abundance after 1 h recovery (E) and 1725 fast-recovery translated mRNAs (F) among the upregulated mRNAs. (G) Venn diagram shows the overlap between fast-recovery transcribed mRNAs and fast-recovery translated mRNAs. (H, I) CTRD, NAD^+^ capping, and circadian TE are significantly overrepresented among the fast-recovery transcribed mRNAs (H), and CTRD and circadian TE are significantly overrepresented among the fast-recovery translated mRNAs (I).

This trend was further supported by differential gene expression analysis using the criteria of pvalue < 0.05 and |log₂FoldChange| > 1. After 1 hour of heat stress at 37 °C, we identified 4,415 differentially expressed genes (DEGs) (Figure 6B). Following 1 hour of recovery at 22 °C, the number of DEGs decreased to 1,398 (Figure 6B), and after 3 hours, only 312 DEGs remained (Figure S5C). A similar pattern was observed for differentially translated genes (DTGs) based on TRAP-seq data, after 1 hour of heat stress at 37 °C, we identified 4,480 DTGs (Figure 6C). Following 1 hour of recovery at 22 °C, the number of DTGs decreased to 1,584 (Figure 6C). These results indicate that transient heat stress at 37°C induces widespread transcriptional and translational alterations in plants, with most affected genes recovering rapidly within 1 hour under normal growth condition. To quantify the impact of heat on translational efficiency, we calculated the ratio of TE of samples after 1 hour of heat stress compared to the TE from control samples without heat stress. Overall, after 1 hour of heat stress at 37 °C, there was a general and rapid decrease in TE (Figure 6D). Interestingly, after 1 hour of recovery at 22 °C, an increased TE was observed (Figure 6D). However, after 3- and 6-hours recovery, the TE decreased. This likely reflects dynamic changes in transcribed mRNAs and translated mRNAs during the recovery process, suggesting that post-transcriptional regulation assists plants in reverting to a normal state after transient heat stress.

To investigate how post-transcriptional regulation helps plants revert to normal states after transient heat stress, we focused on genes whose transcribed mRNA and translated mRNA levels both rapidly returned to normal. We performed an intersection analysis of genes upregulated after 1 hour of heat stress (Recovery_0h_up) and those upregulated at 1 h of recovery (Recovery_1h_up). This analysis identified 1,736 genes specific to Recovery_0h_up, indicating that the transcripts of these genes are fast recovery transcribed mRNAs (Figure 6E). Similarly, we identified 1,725 fast recovery translated mRNAs (Figure 6F). A Venn diagram revealed that 1,170 mRNAs overlapped between the fast recovery transcribed mRNAs and fast recovery translated mRNAs (Figure 6G), suggesting that most genes can return to normal levels at both the transcriptional and translational levels. We next examined the features that facilitate a rapid return to normal at the post-transcriptional regulatory level in plants. We found that CTRD, NAD^+^ capping, and circadian TE were significantly overrepresented among fast recovery transcribed mRNAs (Figure 6H), while CTRD and circadian TE were significantly overrepresented among fast recovery translated mRNAs (Figure 6I). These results indicate that a rapid return of mRNA levels to normal level after heat stress requires the coordinated action of multiple factors.

Furthermore, a total of 2,337 genes were downregulated after 1 hour of heat stress at 37 °C, and 662 were downregulated after 1 hour of recovery. To investigate how post-transcriptional regulation assists downregulated genes in returning to normal levels, we compared the downregulated genes after 1 hour of recovery (Recovery_1h_Down) with those after 1h heat stress (Recovery_0h_Down). This analysis revealed 2,039 genes specific to Recovery_0h_Down (Figure S5E), indicating that the transcripts of these genes rapidly reverted to normal levels within 1 hour of recovery. Similarly, for fast-recovery translated mRNAs, we identified 2,125 genes whose translated mRNAs had reverted to normal levels after 1 hour of recovery (Figure S5F). Totally, 1,136 genes were common between fast recovery transcribed mRNAs and fast-recovery translated mRNAs. A further analysis revealed that CTRD and NAD^+^ capping were significantly underrepresented among fast recovery heat-repressed transcribed mRNAs (Figure S5H), while CTRD was significantly underrepresented among fast recovery heat-repressed translated mRNAs (Figure S5I). These results indicate during the recovery of low abundance mRNAs to normal levels, factors that cause mRNA instability are suppressed, facilitating rapid mRNA accumulation.

Additionally, we found that circadian TE was significantly overrepresented in both upregulated and downregulated mRNAs that rapidly reverted to normal levels (Figure 6H, I and Figure S5H, I). We propose that ribosomes rhythmically modulate translation efficiency in response to mRNA abundance, thereby enabling protein levels to rapidly recover to normal level.

## DISCUSSION

Circadian regulation of gene expression enables plants to anticipate daily environmental changes and optimize physiological processes (*39*). While the transcriptome and translatome have both been shown to exhibit circadian rhythm, the mechanisms driving their coordination remain incompletely understood (*17*). In this study, we systematically dissected the dynamics of transcriptional and translational rhythms in Arabidopsis and revealed that co-translational RNA decay (CTRD), RNA stability factors such as intronless structure and NAD^+^ capping, and translational efficiency (TE) all contribute to shaping the amplitude and timing of rhythmic transcription and translation. In addition, we analyzed the effect of these features on responding to transient heat stress, providing new insights into how biological rhythms adapt to heat challenges.

Our intersection analysis of circadian oscillating transcribed mRNAs and translated mRNAs identified 5,185 genes with rhythmic patterns at both the transcriptional and translational levels (Figure 1B). These genes were classified into four distinct groups based on differences in phase and amplitude (Figure 1C-F). Notably, a substantial subset of genes displayed similar amplitudes but shifted phases, or vice versa, highlighting the post-transcriptional complexity involved in circadian regulation. The S2 group, where amplitude differed between transcribed mRNAs and translated mRNAs without a phase shift, was particularly enriched for genes undergoing CTRD, suggesting that CTRD primarily modulates the amplitude of circadian genes. Our analysis further demonstrated that CTRD-associated genes in Arabidopsis exhibit significantly higher circadian amplitudes. This finding extends previous observations in tomato and supports the notion that CTRD is a conserved mechanism for regulating high-amplitude gene oscillations across plant species (*27*). CTRD genes with higher circadian amplitudes were enriched for photosynthesis-related functions, indicating that CTRD may play a key role in fine-tuning energy-demanding processes in response to diel cues.

In parallel, we further evaluated the contribution of RNA stability features to rhythmic amplitude regulation. Transcripts from intronless genes have been reported to be less stable than those from genes containing introns (19, 33, 34). In this study, we found that Intronless genes were consistently associated with lower oscillation amplitudes across Arabidopsis, tomato, and rice, likely due to their shorter half-lives (Figure 3A-D). Additionally, intronless genes are significantly enriched in the S4 group (Figure 1F), suggesting that the intronless structure may contribute to the asynchrony between translated mRNAs and transcribed mRNAs. Similarly, NAD⁺-capped mRNAs, which are prone to degradation and enriched in S2 group (Figure 1C), may be targeted by CTRD (*35*), exhibited intermediate amplitudes between intronless and CTRD-regulated genes. miRNA-mediated targeting of mRNAs serves as a key component of robust post-transcriptional regulation (*20*). When comparing the amplitudes of 34 miRNA-targeted diurnal genes with other diurnal genes, we found no significant difference (Figure S4). Given the limited number of diurnally expressed miRNA targets available in the current dataset, it remains unclear whether miRNA-mediated mRNA decay contributes to amplitude regulation. Future studies incorporating a broader set of miRNA target transcripts will be needed to address this question. Taken together, our findings indicate that transcript stability and degradation mechanisms, such as CTRD, intronless status, and NAD⁺ capping synergistically influence amplitude modulation within the circadian transcriptome. CTRD-associated genes exhibited the highest amplitudes, intronless genes showed lowest amplitudes, and NAD⁺-capped mRNAs showed intermediate amplitude levels (Figure 4E and F). This result provides new insights into the regulation of rhythmic amplitude.

In addition to transcript stability, we explored how translational efficiency contribute to rhythmic regulation. Our analysis identified 379 genes with circadian fluctuations in TE (Figure 5A), indicating that most genes show strong coupling between transcription and translation. However, genes with circadian TE were significantly overrepresented in the S4 group (Figure 5E), suggesting that TE rhythmicity primarily contributes to phase and amplitude differences between transcribed and translated mRNAs.

During normal growth, plants are frequently exposed to transient heat stress (*40*). However, how post-transcriptional regulation rapidly assists plants in returning to normal states after experiencing transient heat stress remains unclear. We analyzed the impact of transient heat stress on the transcriptome and translatome. One hour exposure to 37 °C induced widespread transcriptional and translational changes (Figure 6B and C). Remarkably, after 1 hour of recovery, most of these changes reverted to normal, suggesting that post-transcriptional regulation plays a key role in balancing RNA abundance under environmental stress. We identified 1,736 upregulated mRNAs and 1,725 upregulated translated mRNAs that returned to normal within 1 hour. Feature analysis revealed that CTRD, NAD^+^ capping, and circadian TE were significantly overrepresented among the 1,736 fast-recovery transcribed mRNAs induced by heat stress, whereas CTRD and circadian TE were significantly enriched among the fast-recovery translated mRNAs (Figure 6E and F). Likewise, circadian TE was significantly overrepresented in downregulated transcribed mRNAs and translated mRNAs that rapidly recovered (Figure S5E and F). Notably, previous studies have shown that CTRD facilitates the rapid degradation of light-induced transcripts during the recovery phase following extreme light exposure (*30*). These findings indicate that the rapid restoration of mRNA levels after extreme light or heat stress requires the coordinated action of multiple post-transcriptional regulatory mechanisms.

In summary, our study reveals that co-translational RNA decay, NAD^+^ capping, intronless structure and translational efficiency collectively shape the amplitude and phase difference of rhythmic transcription and translation in plants. Plants coordinate multiple post-transcriptional mechanisms to restore normal transcribed and translated mRNA abundance after transient heat stress. These findings deepen our understanding of post-transcriptional regulation in circadian systems and provide a foundation for exploring how plants maintain homeostasis under environmental fluctuations.

## MATERIALS AND METHODS

### High-throughput sequencing data sources and preprocessing

The Arabidopsis genome (TAIR10) and gene annotation were downloaded from TAIR (https://www.arabidopsis.org). The raw TRAP-seq and RNA-seq data were downloaded from the GEO database at NCBI (accession number: GSE158444). Raw reads were processed to remove adapters and low-quality sequences. The resulting clean reads were mapped to the Arabidopsis reference genome using STAR with default parameters (version 2.7.11b) (*41*). Gene-level read counts were calculated using HTSeq-count (version 2.0.9) (*42*). To normalize gene expression, RPM (Reads of exon model per Million mapped reads) value was employed. The 3,635 CTRD genes in Arabidopsis were compiled from the studies by Guo et al. (2023) (*26*) and Carpentier et al. (2024) (*31*). The circadian genes of tomato and rice were collected from Zhang et al. (2023) (*27*) and Zhang et al. (2023) (*36*), respectively. The RNA half-life data of wild-type Col-0 was obtained from Carpentier et al.’s study (2024) (*31*). The 5,642 NAD⁺-capped RNAs were collected from Wang et al.’s study (2019) (*22*). The intronless genes were extracted from the genome annotation (GTF) files.

### High-throughput sequencing data processing

Raw reads were processed using Fastp (v0.23.2) for quality control and trimming the adapters (*43*). The filtered reads were then aligned to the Arabidopsis thaliana TAIR10 reference genome with corresponding annotations from Araport11 using STAR (v2.7.11b) with default parameters (*41*). Only uniquely mapped reads were retained for downstream analyses. Differential gene expression (DEG) analysis was performed using the DESeq2 R package with thresholds set at P-value < 0.05 and |log₂(fold change)| > 1 for significance (*44*).

### Identification of circadian genes

The circadian oscillation of transcribed mRNAs, translated mRNAs and TE were identified using R packages “Metacycle2” (*28*). In brief, the key parameters: outIntegration= “onlyIntegration”, timepoints= rep(seq(0, 24, by=3), each=3), cycMethod= “JTK”). A p-value less than 0.01 was considered as the threshold to define circadian genes. The phase and amplitude of each gene were extracted from the output files using Perl scripts. Heatmaps of circadian TE was generated using the R package “pheatmap”, with genes sorted by their peak expression times along the rows.

### Gene Ontology enrichment analysis

The target genes were submitted to PANTHER (http://www.pantherdb.org/) for Gene Ontology (GO) enrichment analysis. The top GO terms were selected for barplot visualization using R packages. Unsupervised clustering was performed using the R package Mfuzz (*45*).

### The analysis for the synchronous and asynchronous rhythmicity of genes with circadian transcription and translation in plants

The circadian genes were identified using R packages “Metacycle2” (*28*). The phase and amplitude values were extracted from the output data. The classification criteria for S1-S4 groups were defined as follows: S1 group: |Δphase| < 1.5 h and |Log_2_(amplitude fold change) | < 0.58; S2 group: |Δphase| < 1.5 h and |Log_2_(amplitude fold change) | ≥ 0.58; S3 group: |Δphase| ≥ 1.5 h and |Log_2_(amplitude fold change) | < 0.58; S4 group: |Δphase| ≥ 1.5 h and |Log_2_(amplitude fold change) | ≥ 0.58.

### Measurement of translational efficiency and identification of genes with circadian translational efficiency

For each gene, gene expression levels are normalized to reads per million (RPM), and TE was calculated as the ratio of TRAP-seq RPM to RNA-seq RPM. Time-series TE data were analyzed using R packages “Metacycle2” (*28*), with a pvalue less than 0.01 was considered as the threshold to define circadian TE.

### Unsupervised cluster

The unsupervised cluster was conducted using the Mfuzz R package (*45*). The RPM values of total mRNAs and TRAP mRNAs as input date. In brief, the key parameters: mfuzz.plot2(data, cl, mfrow=c(1,4), time.labels=c(”ZT48”, “ZT51”, “ZT54”, “ZT57”, “ZT60”, “ZT63”, “ZT66”, “ZT69”, “ZT72”), min. mem=0.5, centre=TRUE, centre.col=c(”deepskyblue”), centre.lwd=2, ylab=”Scaled Normalized expression values”, x11=F, colours = brewer.pal(n = 4, name = “Greys”)).

### Code availability

All the scripts used in this study were written by the author, and are available upon request to the corresponding author.

## ACKNOWLEDGEMENTS

This work was funded by grants from National Natural Science Foundation of China (Grant No. 32170581 and No. 32370587 to XY), Shanghai Municipal Commission of Education (Grant No. 2024AIYB005), and this research is funded by the Hong Kong Scholars Program.

## AUTHOR CONTRIBUTIONS

PX and XY designed the study. PX, YW, QW compiled all sequencing data and carried out the computational analysis. PX and XY wrote the manuscript. All authors read and approved the manuscript.

## SUPPORTING INFORMATION

The following materials are available in the online version of this article.

Table S1. The list of circadian transcripts in RNA-seq and TRAP-seq datasets.

Table S2. The list of CTRD genes used in this study.

Figure S1. GO enrichment analysis of 5,185 circadian common genes.

Figure S2. CTRD-associated circadian genes are enriched in photosynthesis pathways and exhibit the highest amplitude oscillations.

Figure S3. Intronless genes exhibit low circadian amplitude in Arabidopsis and rice. (A) The frequency of intronless genes in the whole genome. (B, C) Coding genes with or without introns display significantly different circadian amplitudes in Arabidopsis root (B) and rice (C). (D) Intronless genes exhibit low circadian amplitude in translated mRNAs

Figure S4. MiRNAs do not contribute to amplitude variation. (A) Venn diagram showing the overlap among circadian total mRNAs and miRNA-targets. (B) There is no significant difference in amplitude between miRNA target genes and non-target genes.

Figure S5. Plants coordinate CTRD and circadian TE to restore genes suppressed by heat stress to normal levels. (A, B) Principal component analysis of transcribed mRNAs (A) and translated mRNAs (B), based on RPM-normalized expression values. The circle highlights the outlier population. (C, D) Volcano plot showing differentially expressed genes after 3-6 hours recovery. The identified DEGs in RNA-seq (C) and DTGs in TRAP-seq (D). (E-G) Venn diagram analysis of fast recovery transcribed mRNAs and fast recovery translated mRNAs among downregulated mRNAs. (H, I) Among the downregulated mRNAs, CTRD was significantly underrepresented in both the fast-recovering transcribed mRNAs (H) and fast-recovering translated mRNAs (I).

